# Glia-Mediated Antigen Presentation In The Retina During Degeneration

**DOI:** 10.1101/2024.04.21.590440

**Authors:** Simona Intonti, Despina Kokona, Martin S. Zinkernagel, Jens V. Stein, Volker Enzmann, Federica M. Conedera

## Abstract

Glia antigen-presenting cells (APCs) are pivotal regulators of immune surveillance within the retina, maintaining tissue homeostasis and promptly responding to insults. The intricate mechanisms underlying their local coordination and activation remain unclear.

Our study integrates an animal model of retinal injury, retrospective analysis of human retinas, and in vitro experiments to elucidate insights into the pivotal role of antigen presentation in neuroimmunology during retinal degeneration, uncovering the involvement of various glial cells, notably Müller glia, and microglia. Glial cells act as sentinels, detecting antigens released during degeneration and interacting with T-cells via MHC molecules, which are essential for immune responses. Microglia function as APCs via the MHC class II pathway, upregulating key molecules such as Csf1r and cytokines. In contrast, Müller cells act as atypical APCs through the MHC class I pathway, exhibiting upregulated antigen processing genes and promoting a CD8^+^ T-cell response. Distinct cytokine signaling pathways, including TNF-α and IFN, contribute to the immune balance. Human retinal specimens corroborate these findings, demonstrating glial activation and MHC expression correlating with degenerative changes. In vitro assays also confirmed differential T-cell migration responses to activated microglia and Müller cells, highlighting their role in shaping the immune milieu within the retina. These insights emphasize the complex interplay between glial cells and T-cells, influencing the inflammatory environment and potentially modulating degenerative processes.

In summary, our study emphasizes the involvement of retinal glial cells in modulating the immune response after insults to the retinal parenchyma. Thus, unraveling the intricacies of glia-mediated antigen presentation in retinal degeneration is essential for developing precise therapeutic interventions for retinal pathologies.

## INTRODUCTION

The eye was once thought to be an immune privileged site; however, this notion is increasingly challenged by evidence of T-cell infiltration in many retinal diseases (Camelo, 2014). This prompts further questions into the role of the immune system in the development of retinal diseases. While the role of T-cells in diseases like uveitis is well established, others as age-related macular degeneration (AMD), are poorly explored to date. We provided significant evidence of T-cell accumulation in ocular tissues of humans and demonstrated the harmful effect of cytotoxic (CD8^+^) T-cells on retinal degeneration in a murine model of focal injury (Conedera et al., 2023). While these data suggest T-cells are implicated in retinal degeneration, the precise mechanisms of their activation and interaction remain elusive, indicating a need to explore how antigen-presenting cells (APCs) orchestrate adaptive immune responses within the retina.

The retina harbors two subsets of APCs known as microglia and macroglia. Both glia cell types play pivotal roles in immune responses by actively surveying their microenvironment, presenting antigens to T-cells, and orchestrating immune tolerance (Reichenbach & Bringmann, 2020). Microglia, the resident immune cells of the retina (Murenu et al., 2022), constitute 5-20% of the total glial cell population within the retinal parenchyma (Conedera et al., 2019). Under physiological conditions, they exhibit a ramified morphology and continuously scan the retinal parenchyma using motile protrusions, playing a vital role in immune surveillance (Verkhratsky et al., 2021). These processes enable microglia to promptly detect abnormalities within retinal tissue, facilitating rapid responses to potential threats. During degeneration, microglia become activated in response to stimuli such as inflammation or tissue damage. Consequently, they release factors and express receptors to modulate immune responses and tissue repair (Damani et al., 2011). Upon activation, microglia prime and activate autoreactive T-cells by upregulating major histocompatibility complex (MHC) molecules and expressing co-stimulatory molecules like CD80 and CD86 (Guo et al., 2022; Jurga et al., 2020). Additionally, microglia release cytokines that attract T-cells to the retinal parenchyma, thereby contributing to immune responses in retinal degenerative diseases (Ramirez et al., 2017). Müller cells span the entire depth of the retina, contributing to light transmission and maintaining homeostasis under physiological conditions (Szabó et al., 2022; Yoshimoto et al., 2023). In response to retinal degeneration, they exhibit reactivity to protect the retina from further damage and promote repair following pathological insult. Reactive gliosis includes morphological, biochemical and physiological changes, which vary depending on the type and degree of the injury (Graca et al., 2018). Reactive Müller cells are characterized by an altered expression of both non-specific and specific markers, such as glial fibrillary acidic protein (GFAP) and glutamine synthetase (GS), indicating their responsiveness to retinal diseases and injuries (Bringmann et al., 2006). Simultaneously, Müller cells effectively express both MHC class I and II molecules enabling them to actively interact with CD4^+^ and CD8^+^ T-cells (Lorenz et al., 2021).

Thus, microglia and Müller cells can potentially interact with different subsets of T-cells through the expression of MHC molecules during retinal degenerative diseases. However, how glial cells coordinate the T-cell response to retina damage remains unknown. There is a compelling need to understand the specific role of glial cells in T-cell modulation during retinal degeneration and how their interaction governs complex cell–cell interactions that, in turn, may influence the retina’s ability to recover from injury.

Here, we combined an animal model of retinal injury, retrospective studies on human retinas and *in vitro* experiments to determine the role of glial cells on the T-cell response to retinal damage. Such information is fundamental to understanding retinal disorders and developing targeted therapeutic strategies that can safely modulate the immune response to retinal degeneration and improve outcomes.

## METHOD

### Animals

All animal experiments were approved by the local Animal Ethics Committee of the Canton of Bern (Switzerland; BE34/19) and conform to the Association for Research in Vision and Ophthalmology Statement for the Use of Animals in Ophthalmic and Vision Research. Male and female 8-12 weeks old C57Bl/6J mice were purchased from Charles River (Sulzfeld, Germany). B6-Tg (Rlbp1^GFP^) mice (4–8 weeks old) were originally provided by Prof. Dr. Christian Grimm. Genotyping of Rlbp1^GFP^ mice was performed as previously described (Graca et al., 2018). Csf1r^EGFP^ mice were acquired from Jackson Laboratory (Bar Harbor, ME, USA; Strain #005070) and express enhanced green fluorescent protein in resident macrophages and microglia in the retina. All animals were housed in designated animal holding facilities observing a standard twelve-hour day/night cycle. Standard rodent chow and water were provided ad libitum.

### Retinal laser-injury

Focal injury to the retina was induced as previously described (Conedera et al., 2021). Mice were anesthetized by subcutaneously injecting 45 mg/kg ketamine (Ketalar 50 mg/ml; Pfizer AG, Zurich, Switzerland) and 0.75 mg/kg medetomidine hydrochloride (Domitor, 1 mg/ml; Orion Pharma AG, Zug, Switzerland). Six lesions were created on both eyes using a 532 nm diode laser (Visulas 532 s, Carl Zeiss Meditec AG, Oberkochen, Germany). For the RNAseq analysis, we induced retinal damage by generating up to 50 laser burns per eye, encompassing the visible portion of the fundus. Each burn was produced with 120 mW of power for 60 ms and 100 μm in diameter.

### Retinal dissociation, sorting, and RNA-Seq library production

Retinas of Csf1r^EGFP^ or Rlbp1^GFP^ mice (n = 3) were dissected at days 1, 3, and 7, and digested with papain (Worthington Biochemical, Freehold, NJ, USA) for 15 min (Feodorova et al., 2015). After dissociation, cell suspension was incubated in HBSS with 0.4 % BSA (ThermoFisher Scientific, Basel, Switzerland) and DNase I (200 U/ml; Sigma-Aldrich, Buchs, Switzerland) and filtered with a 35 μm cell strainer (Costar Corning, Cambridge, MA, USA). To evaluate cell viability, we supplemented added Hoechst 33342 Ready Flow™ Reagent (ThermoFisher Scientific) to the cell suspension. Background fluorescence was determined using cells from Csf1r^EGFP^- or Rlbp1^GFP^-negative littermates.

We sorted 100 Csf1r^EGFP^- or Rlbp1^GFP^-positive cells/μl using Moflo Astrias (Beckman-Coulter, Nyon, Switzerland) into Buffer TCL (4 μl; Qiagen, Venlo, Netherlands) containing 1 % 2-mercaptoethanol (Sigma-Aldrich). After cell sorting, all samples were processed using the published Smart-seq2 protocol4 to generate the cDNA libraries (Picelli et al., 2014). Subsequently, the libraries were sequenced on an Illumina HiSeq4000 (Illumina, San Diego, CA, USA) with a depth of approximately 20 million reads per sample. Sequencing data are available as an additional files.

### RNA sequencing

The raw sequencing reads underwent initial processing, including the removal of adapter sequences, trimming of low-quality ends, and the exclusion of reads with a phred quality score below 20, achieved through the utilization of Trimmomatic (Version 0.36). Subsequent read alignment was executed utilizing STAR (v2.6.0c). The Ensembl murine genome build GRCm38.p5, augmented with gene annotations retrieved on 2018-02-26 from Ensembl (release 91), served as the reference genome for the alignment process. The STAR alignment options were “--outFilterType BySJout --outFilterMatchNmin 30 --outFilterMismatchNmax 10 --outFilterMismatchNoverLmax 0.05 -- alignSJDBoverhangMin 1 --alignSJoverhangMin 8 --alignIntronMax 1000000 --alignMatesGapMax 1000000 -- outFilterMultimapNmax 50”. Gene expression values were computed with the function featureCounts from the R package Rsubread (v1.26.0). The options for feature counts were: - min mapping quality 10 - min feature overlap 10 bp - count multi-mapping reads - count only primary alignments - count reads also if they overlap multiple genes. To identify genes showing differential expression, we employed a count-based negative binomial model as implemented in the software package DESeq2 (R version: 3.5.0, DESeq2 version: 1.20.0). The assessment of differential expression employed an exact test tailored for over-dispersed data. Genes displaying altered expression with an adjusted p-value < 0.05 (Benjamini and Hochberg method) were designated as differentially expressed. Subsequently, heatmaps were constructed for specific gene subsets using the heatmap.2 function from the gplots package (v. 3.0.1) in R v. 3.5.1. These heatmaps visually represented the log2 fold-changes between the two experimental groups.

### Human specimen and tissue processing

Retrospective studies were performed on post-mortem eyes obtained from donors’ eyes after the removal of the cornea for transplantation. Within 24 h post-mortem, retinal samples were fixed with 4 % formaldehyde at 4°C overnight, embedded in paraffin and subjected to pre-established inclusion and exclusion criteria. We included retinas from patients of both sexes older than 60-year-old, as retinal degenerative diseases usually begin at that age. Eyes from donors suffering from systemic or ocular comorbidities were excluded based on the criteria for the corneal donations (e.g., sepsis, meningitis, HIV, lues, hematological neoplasms, all ocular tumors, Creutzfeldt– Jakob disease, rapid progressive dementia or degenerative neurological condition, eye surgery within 6 months or after transplantations, drug abuses). The research was approved by the Ethics Committee of the Canton of Bern (2022-01842). Paraffin sections (5 μm) were stained with Mayer’s hemalum and eosin (H&E; Roth, Karlsruhe, Germany) to evaluate the presence of hyalinized deposits between the retinal pigment epithelium (RPE) and Bruch’s membrane and define the number of nuclei per retina, as previously reported (Gupta et al., 2016; Li et al., 2018; Quinn et al., 2019). Sections were deparaffinized, rehydrated and immersed in a Mayer’s hemalum solution for 5 minutes. Following this, the slides were immersed in an eosin dye (Sigma-Aldrich). Finally, we briefly rehydrated the sections before mounting them using a mounting medium (Vector Laboratories, Burlingame, CA, USA). Additionally, we stained retinal samples with Picro Sirius red staining specifically designed for histological visualization of collagen fibers and thus tissue fibrosis. Paraffin sections underwent deparaffinization and hydration. Subsequently, the nuclei were stained using Weigert’s hematoxylin (Roth), for 5 minutes, followed by rinsing with tap water for 10 minutes. After, sections were incubated in Direct Red 80 (Sigma-Aldrich) for 1 hour and 30 minutes allowing it to interact with the collagen fibers. Subsequently, stained samples were washed with acidified water, dehydrated and mounted using the mounting medium for microscopic examination.

High-throughput and high-quality brightfield H&E- and Sirius Red-stained images of the human retina were acquired with a NanoZoomer 2.0-HT slide scanner (Hamamatsu Photonics France, Massy, France). We examined retinas from 30 donors, which were subdivided in two groups (n = 15 per group): retinas not showing pathological features of retinal degeneration (control, CTRL) and 15 presenting drusen (> 25 μm), retina atrophy (< average number of cell nuclei) and fibrosis (retinal degeneration, RD).

### Immunofluorescence

Paraffin sections (5 μm) from human and murine eyes were utilized for immunofluorescence. Antigen retrieval was achieved by incubating the sections in either Tris–EDTA (pH 9.0) or citrate buffer (pH 6.0) with 0.05 % Tween-20 for 20 min, then cooling at room temperature for 30 min. Sections were blocked for 1 h in Tris-buffered saline + 5 % goat normal serum + 1 % bovine serum albumin (pH 7.6) and incubated with primary antibodies overnight at 4°C. Primary antibodies used in this study were: rabbit anti-ionized calcium-binding adapter molecule 1 (Iba1; 1:500; 019-19741, Wako Pure Chemical Industries Ltd., Osaka, Japan), rabbit anti-glutamine synthetase (GS; 1:200; ab197024, Abcam, Cambridge, UK), mouse anti-major histocompatibility complex class I (MHC I; 1:200; 311402, BioLegend, San Diego, CA, USA), mouse anti-major histocompatibility complex class II (MHC II; 1:200; 327002, BioLegend), mouse anti-inducible nitric oxide synthase (iNOS; 1:200; MA5-17139, Invitrogen, Carlsbad, CA, USA), and mouse anti-glial fibrillary acidic protein (GFAP; 1:200; OPA1-06100, Invitrogen). This step was followed by washing with PBS and 0.05 % Triton X-100 and incubation with the respective secondary antibodies conjugated to 488/594 fluorophores (Alexa Fluor® 488 and Alexa Fluor® 594, Abcam). Cell nuclei were counterstained using the mounting media Vectashield with DAPI (Vector Laboratories, Newark, CA, USA).

### Cell lines

The human microglial clone 3 cell line (HMC-3), was purchased from ATCC (CRL-3304; Manassas, VA, USA). HMC-3 cells were cultured in Dulbecco’s minimum essential medium (DMEM; Gibco, Scientific, MA, USA) supplemented with GlutaMAX (Gibco), 10% fetal bovine serum (FBS; Gibco) and 1% antibiotic/antimycotic (A/A; Gibco). The cells were maintained in a humidified incubator at 37°C and 5% CO_2_ and were promptly split as soon as they reached confluency. The spontaneously immortalized MIO-M1 cell line was provided by G.A. Limb (Limb et al., 2002). This cell line was cultured in DMEM, to which 10% FBS and 1% A/A was added. MIO-M1 cells were cultured at 37° C and 5% CO_2_.

### Isolation of human primary T-cells

Primary human T-cells were isolated from anonymized, donated buffy coats from the Interregional Blood Bank Bern (Project P_406). The separation was performed by centrifugation via a Ficoll-Paque (Gibco) density gradient. To begin the isolation process, the buffy coat was diluted with Dulbecco’s Phosphate-Buffered Saline (DPBS; Gibco) in a 1:1 ratio in a conical tube (Corning, New York, NY, USA) and mixed gently. Subsequently, the diluted buffy coat was carefully added onto the Ficoll-Plaque, ensuring that the two phases are not mixed, and centrifuged at 900g. Following centrifugation, the PBMC layer is transferred into a separate tube containing 10 ml of DPBS, mixed gently, and centrifuged at 250g. After centrifugation, the supernatant is discarded, and the pellet, containing the cells, is resuspended in Mojo sorting buffer (BioLegend) composed of DPBS supplemented with 1 % A/A (Gibco) and 2.5 % FBS. Cell count assured that the cell number falls within the range of 1x10^7^ to 2x10^8^ for using the MojoSort™ human CD3 T-cell isolation kit (480022; BioLegend). The procedure was performed as recommended by the manufacturer. In brief, 10 μL of the biotin-antibody cocktail was added to the filtered cells. The sample is mixed gently and placed on ice for 15 minutes. Next, 10 μL of streptavidin nanospheres are added to the sample and left on ice for an additional 15 minutes. Afterward, the sample was placed inside the magnet for 5 minutes. The CD3^+^ T-cells, which have bound to the streptavidin nanospheres, were then harvested for further analysis. CD3^+^ T-cells were plated in 24-well plates (1 x 10^7^ cells per well) and cultured in RPMI medium supplemented with 10 % FBS and 1 % A/A. To activate CD3^+^ T-cells, we stimulated them with a human anti-cluster of differentiation 3 antibody (CD3; 1:200; 300465, BioLegend) and then incubated at 37°C for 72 hours.

### *In vitro* migration assay

To determine the migration of T-cell toward APCs, a transwell assay was performed. To evaluate the effect of activated T-cells on HMC-3 and MIO-M1, a 12-well tissue culture insert with a pore size of 8.0 μm was utilized (Corning). Therefore, the HMC-3 cells were detached using trypsin (Gibco), centrifuged, and re-suspended in RPMI medium supplemented with GlutaMAX, 10 % FBS, 1 % A/A. The same procedure was performed for MIO-M1, but they were suspended in RPMI medium containing 10 % FBS and 1 % A/A. Then, HMC-3 and MIO-M1 cells were seeded in the lower chamber (4x 10^4^ cells per well). In the upper chambers, T-cells (1x10^6^ per well) were placed. After 3 hours, the migrated T-cells were collected, and immunocytochemical staining was conducted using rabbit anti-CD4 (1:200; ab133616, Abcam) and mouse anti-CD8 antibodies (1:100; ab17147, BioLegend). This procedure was performed in triplicate.

### Activation of microglia

When the HMC-3 cells reached confluency, they were detached using trypsin at 37°C for 5 min, centrifuged at 300g for 5 min, and re-suspended in RPMI. Subsequently, HMC-3 cells were seeded in 8-well chambers (Corning, New York, NY, USA) (4 x 10^4^ cells per well), incubated overnight to adhere, and starved in RPMI without FBS for 6 h. Afterward, HMC-3 cells were treated with LPS (2 μg/ml; Sigma-Aldrich) for 24 h to reach a pro-inflammatory state. This procedure was performed in triplicate.

### Treatment of glia cells with either T-cell-derived media or activated T-cells

After activation, T-cells were centrifuged at 300g to separate the supernatant from the cell pellet. The latter, consisting of the activated T-cells, was then gently resuspended in RPMI medium supplemented with 10 % FBS and 1 % A/A. In the next step, HMC-3 or MIO-M 1 cells were incubated with the resuspended pellet and the separated supernatants separately.

### Immunocytochemistry

To examine the cellular interactions, immunocytochemistry was performed in an 8-well chamber slide (Corning) containing HMC-3 or MIO-M1 cells in contact with T-cells and T-cell-derived media, respectively. This approach allowed for the visualization and analysis of specific markers, as mouse anti-iNOS (1:200; MA5-17139, Invitrogen) and mouse anti-GFAP (1:200; OPA1-06100, Invitrogen). To perform this staining, the cells were fixed with 4 % paraformaldehyde (Sigma-Aldrich) at room temperature for 30 minutes. After DPBS washing, the cells were blocked with 5 % normal goat serum (Dako, Glostrup, Denmark) in DPBS supplemented with 0.5 % Triton-X-100 to minimize non-specific binding for 30 minutes. Primary antibodies were added in DPBS, and the cells were incubated at 4°C overnight. The wells were washed three times with DPBS, followed by incubation with the secondary antibodies, goat anti-rabbit/anti-mouse Alexa 488/594 (1:500; Invitrogen) diluted in DPBS at room temperature for 1 hour. The nuclei were counterstained with DAPI (1:15000; Sigma-Aldrich) at room temperature for 5 minutes, and the wells were washed again before being mounted with Vectashield mounting medium (Vector Laboratories). The cells were then imaged using a confocal microscope (LCI Zeiss LSM 710), and the percentage of positive cells for each marker was calculated using ImageJ software (NIH, Bethesda, MD, USA).

### Statistical analysis

Statistical analysis was performed using GraphPad Prism (version 7.0, GraphPad Software, La Jolla, USA). Intergroup comparisons were based on a non-parametric one-way analysis of variance (ANOVA) and the Bonferroni multiple comparison post hoc test. All results are expressed as the mean ± standard deviation (SD). The level for statistical significance was set at a p value ≤ 0.05.

## RESULTS

### Resident immune cells act as APCs in response to injury in the retina via the MHC Class II pathway

Microglia and resident macrophages represent the main APC inside the retinal parenchyma (Schetters et al., 2017). The contribution of APCs in a disease like uveitis is well established, while the role of T-cells in degenerative diseases, such as AMD, is poorly understood (Sutter & Crocker, 2022). We utilized a focal laser injury model that mimics key features of AMD, including photoreceptor degeneration, local inflammatory responses, leukocyte recruitment, as well as glial reactivity and scarring, mirroring observations in damaged retinas in humans (Conedera et al., 2021).

We assessed the expression of MHC molecules in resident phagocytic cells (Iba1^+^) as a key characteristic of APCs (Fig. 1 a-d). Before injury and at 24 hours post-injury, Iba1^+^ cells showed negative staining for both classes of MHC molecules (Fig. 1 b, d). However, starting from day 3 post-injury, we observed an increase in their protein levels (Fig. 1 a-d). Although a 9% increase in MHC I expression was detected in Iba1^+^ cells, it did not significantly differ from baseline (p-value = 0.15, Fig. 1 a-b). By day 7, only 2% of resident phagocytic cells expressed MHC I, similar to the level found before injury (Fig. 1 b). In contrast, MHC II protein levels significantly increased compared to baseline and day 1 on days 3 and 7 post-injury (Fig. 1 c-d). We found that 60% of Iba1^+^ cells were MHC II positive on day 3, with only 36% remaining positive on day 7; however, there was no statistical difference between these two time points (Fig. 1 c-d).

**Fig. 1:**
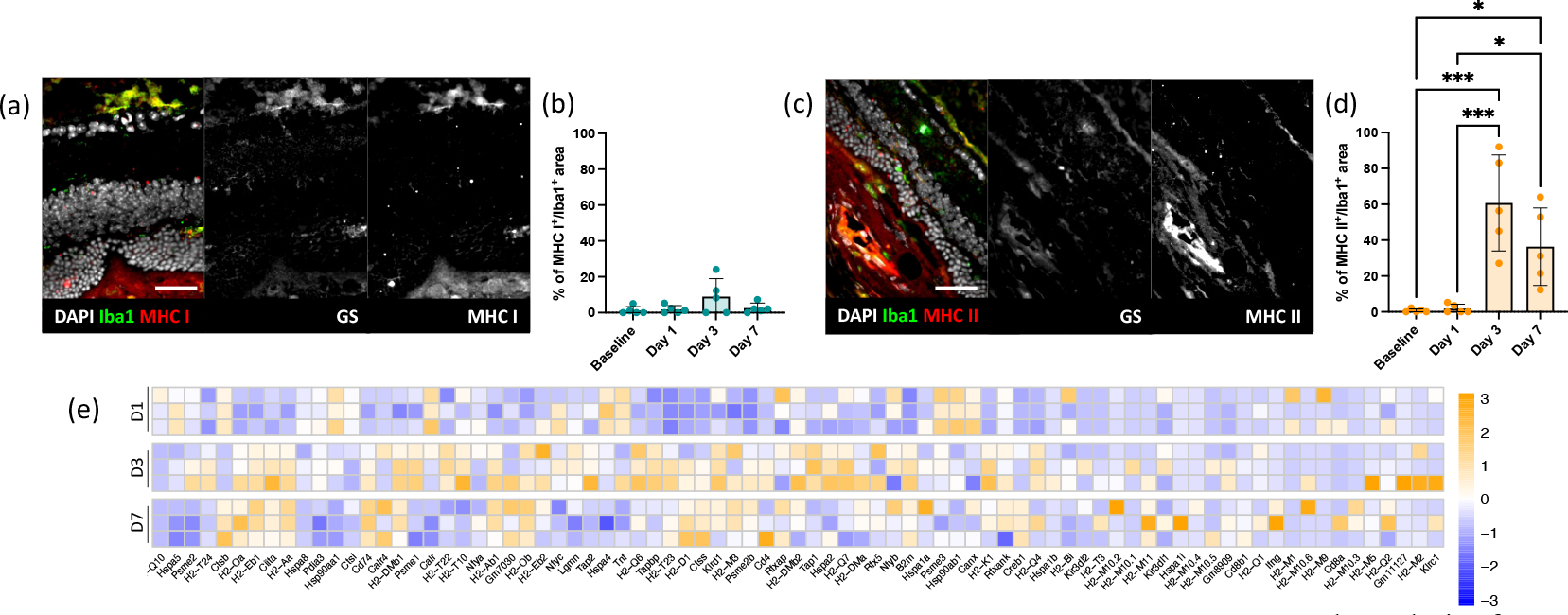
Retinal injury triggers microglial MHC II expression and CD4 T-cell signaling upregulation. (a-d) Analysis of MHC proteins by microglia on day 3 after injury using a microglia marker (Iba1) and MHC class I marker (MHC I). (a) Shown are representative sections stained for Iba1 (green) and MHC I (red). Scale bars equals 100 μm. (b) Quantification of the MHC I^+^Iba1^+^ area on total Iba1^+^ area per lesion before injury (baseline) and at pre-defined time points (days 1, 3 and 7). Significant differences between baseline and the different time points were determined by using a post hoc Bonferroni one-way ANOVA test (n = 5). (c) Shown are representative sections stained for Iba1 (green) and MHC II (red). Scale bars equals 100 μm. (d) Quantification of the percentages of the MHC II^+^Iba1^+^ area on total Iba1^+^ area per lesion before injury (baseline) and at pre-defined time points (days 1, 3 and 7). Significant differences (*p < 0.1 and ***p < 0.001) between baseline and the different time points were determined by using a post hoc Bonferroni one-way ANOVA test (n = 5). (e) Heatmaps of differentially expressed APC-related genes in Csfr1^EGFP^ cells. Genes were selected from KEGG pathways (mmu04612). Data are expressed as fold-changes between different time points (days 1, 3, and 7) compared to negative controls (Csfr1^EGFP^ cells from uninjured retinas).

To validate our immunofluorescence data, we conducted a transcriptome analysis of phagocytic cells expressing the colony-stimulating factor 1 receptor (Csf1r) before and after injury at specified time points (days 1, 3, and 7). Using RNAseq from Csf1r^+^ cells, we identified more than 90 genes related to antigen processing and presentation with significant fold-changes (p-value < 0.05; Fig. 1 e, S1). Interestingly, the expression of these genes was predominantly upregulated on days 3 and 7 post-injury concomitant with the MHC II expression in IBA1^+^ cells (Fig. 1 e). Among the significantly upregulated transcripts, we observed histocompatibility 2 class II antigen E beta2 (H2-Eb2), HLA class II histocompatibility antigen gamma chain (Cd74) and class II major histocompatibility complex transactivator (Ciita; Fig. e, S1). These genes have previously been shown to play crucial roles in the MHC II pathway, contributing to the antigen presentation process essential for adaptive immune responses (Accolla et al., 2019; Cloutier et al., 2021; Logunova et al., 2023). In particular, H2-Eb2 facilitates the presentation of peptides to CD4^+^ T-cells, while Cd74 aids in stabilizing and directing newly synthesized MHC II molecules. Ciita regulates MHC II gene transcription by binding to their promoters and recruiting transcriptional machinery, thereby essential for the expression of MHC II molecules on the surface of antigen-presenting cells.

These data indicate that resident phagocytic cells upregulated MHC II and its pathway, suggesting their ability to present antigens to CD4^+^ helper T-cells.

Furthermore, our RNA sequencing data revealed an upregulation of certain cytokines and pro-inflammatory molecules known to enhance the expression of MHC molecules during the injury response 24 hours after injury (Fig. 2 a-c, S2). Csf1r, however, was significantly upregulated on days 3 and 7 compared to baseline (Fig. 2 a, S2). These findings provide evidence that CSF signaling may promote immune cell differentiation and activation, potentially influencing their capacity to effectively present antigens via the MHC II pathway. Cytokines such as interferons (IFNs) and tumor necrosis factor alpha (TNF-α) are known to enhance the expression of MHC molecules, thereby facilitating antigen presentation and immune responses during inflammation or immune activation. However, we did not find a statistically significant difference in the expression of either interferons or TNF-α pathway genes compared to basal levels.

**Fig. 2:**
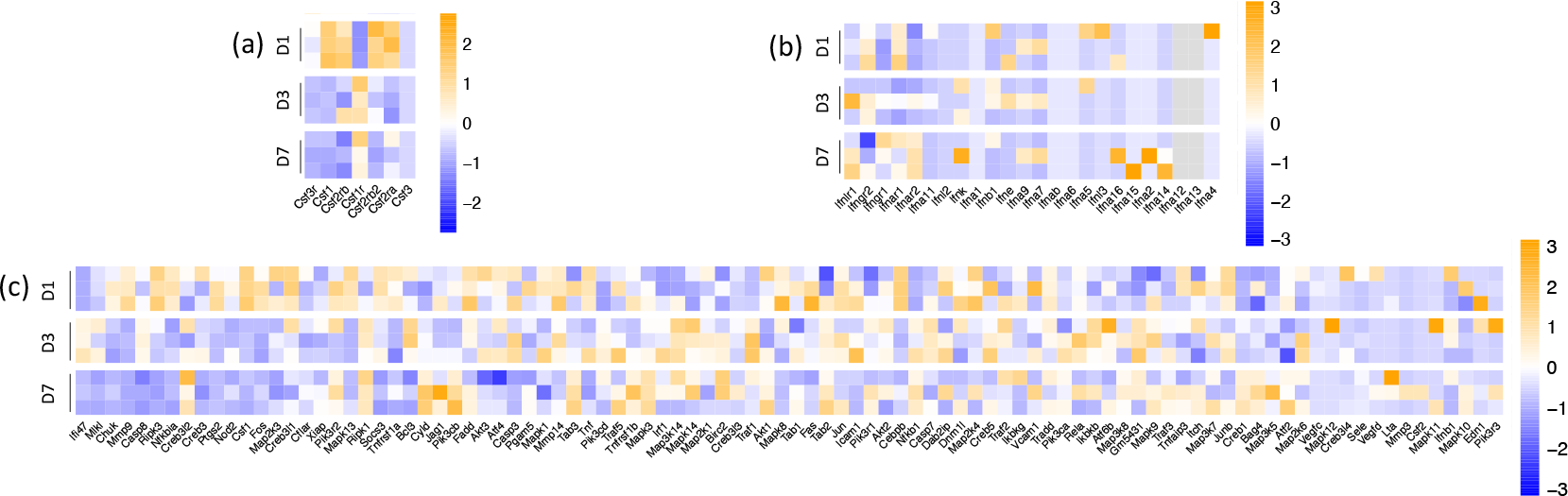
Cytokines release by microglia cells in response to laser injury. (a) Heatmaps of differentially expressed genes that encode receptors for CSFs in Csfr1^EGFP^ (microglia). Data are expressed as fold-changes between different time points (days 1, 3, and 7) compared to negative controls (Csfr1^EGFP^ cells from uninjured retinas). (b) Heatmaps of differentially interferon receptor-ligand genes in Csfr1^EGFP^ (microglia). Data are expressed as fold-changes between different time points (days 1, 3, and 7) compared to negative controls (Csfr1^EGFP^ cells from uninjured retinas). (c) Heatmaps of differentially expressed genes of the TNF signaling pathway in Csfr1^EGFP^ (microglia). Genes were selected from KEGG pathways (mmu04668). Data are expressed as fold-changes between different time points (days 1, 3, and 7) compared to negative controls (Csfr1^EGFP^ cells from uninjured retinas).

These data demonstrate an enhanced activation and differentiation of resident phagocytic cells, potentially influencing their ability to effectively present antigens. Moreover, we found that interferons and TNF-α may not play primary roles as mediators in this context.

### Müller cells are atypical APCs via the MHC Class I pathway in the injured retina

Apart from microglia, Müller cells showed several characteristics of APCs, as they are capable of inducing both MHC class I and MHC class II with co-stimulatory molecules *in vitro* (Schmalen et al., 2021). However, there is limited literature on their APC ability in response to retinal damage in murine models.

We assessed the expression of MHC molecules in Müller glia (GS^+^) in response to focal injury to the retinal parenchyma (Fig. 2 a-d). Similar to our observations in microglia, GS^+^ cells were negative for both classes of MHC molecules before injury and at 24 hours post-injury (Fig. 2 b, d). We observed that on day 3 post-injury, MHC I protein levels in Müller cells showed a slight increase, with 8% positivity. However, this increase only reached statistical significance on day 7 post-injury, when over half of GS^+^ cells expressed MHC I (62%). In contrast, MHC II protein levels in Müller cells remained similar to basal levels throughout the injury response, showing a non-significant increase on day 7 post-injury compared to baseline (p-value = 0.27, Fig. 2 c-d).

To investigate whether Müller cells actively participate in the injury response as APCs, we conducted a transcriptome analysis of retinaldehyde-binding protein 1 (Rlbp1)-positive cells. In the context of retinal injury or disease, Müller cell proliferation often occurs in response to damage as part of the retinal repair process, potentially leading to the formation of a glial scar. Thus, we analyzed only Rlbp1^+^ cells that re-enter the cell cycle. RNA sequencing was performed in reactive Müller cells before and after injury at specified time points (days 1, 3, and 7). We observed that most of the genes related to antigen processing and presentation were predominantly upregulated 7 days after injury, coinciding with the expression of MHC I in GS^+^ cells (Fig. 3 e). Among the significantly upregulated transcripts, we observed proteasome activator subunit 1 (Psme1), transporter 2 ATP binding cassette subfamily b member (Tap2), TAP binding protein (Tapbp) and calnexin (Canx; Fig. 3 e, S3). Psme1 assists in the generation of peptides for MHC class I presentation, Tap genes facilitate the transport of peptides for MHC class I loading, and Canx aids in the folding and assembly of MHC class I molecules. These genes collectively play important roles in the MHC class I antigen processing and presentation pathway, contributing to the immune response by presenting intracellular antigens to cytotoxic T-cells. These data indicate that Müller glia are atypical APCs upregulating MHC I and its pathway and presenting antigens to CD8^+^ cytotoxic T-cells.

**Fig. 3:**
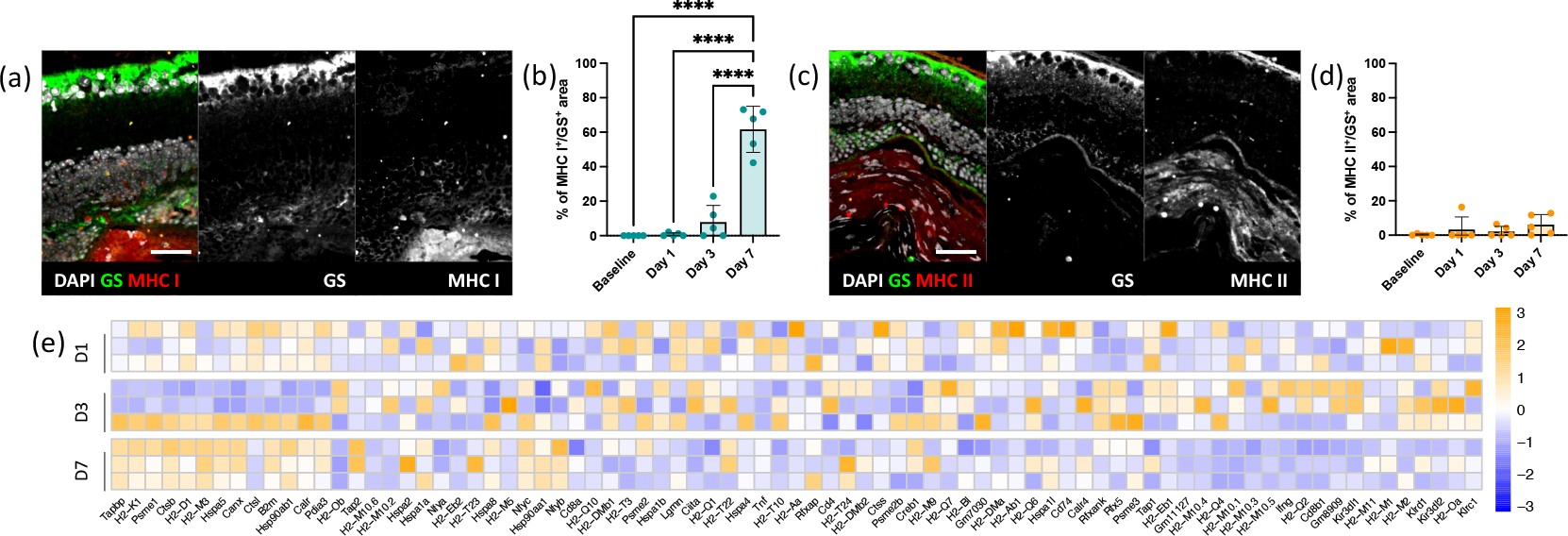
Retinal injury induces the expression of MHC class I in Müller glia and upregulates CD8 T-cell signaling. (a-d) Analysis of MHC proteins expression by Müller cells on day 7 after injury using a Müller cell marker (GS) and MHC class I marker (MHC I). (a) Shown are representative sections stained for GS (green) and MHC I (red). (b) Quantification of the percentages of the MHC I^+^GS^+^ area on total GS^+^ area per lesion before injury (baseline) and at pre-defined time points (days 1, 3 and 7). Significant differences (****p < 0.0001) between baseline and the different time points were determined by using a post hoc Bonferroni one-way ANOVA test (n = 5). Scale bars equals 100 μm. (c) Shown are representative sections stained for GS (green) and MHC II (red). (d) Quantification of the percentages of the MHC II^+^GS^+^ area on total GS^+^ area per lesion before injury (baseline) and at pre-defined time points (days 1, 3 and 7). Significant differences between baseline and the different time points were determined by using a post hoc Bonferroni one-way ANOVA test (n = 5). Scale bars equals 100 μm. (e) Heatmaps of differentially expressed APC-related genes in Rlbp1^GFP^ cells. Genes were selected from KEGG pathways (mmu04612). Data are expressed as fold-changes between different time points (days 1, 3, and 7) compared to negative controls (Rlbp1^GFP^ cells from uninjured retinas).

Additionally, our RNA sequencing data showed an elevation in certain signaling molecules associated with inflammation, known for their capacity to act on MHC pathways during injury response (Fig. 4 a-c, S4). Müller cells have an increased gene expression of CSF signaling similar to what observed in the resident immune cell response. However, reactive macroglia upregulated only Csf2ra 7 days post injury, rather than Csf1r (Fig. 4 a, S4). The expression of Csf2ra in Müller cells may be connected to their immunomodulatory functions and their potential role in antigen presentation via MHC I molecules, contributing to immune surveillance and responses within the retina. Interestingly, we found significant differences in IFN expressions, especially Infar1 and 2, compared to their basal level (Fig. 4 a, S4). Activation of Infar1 and Infar2 initiates a signaling cascade that leads to the upregulation of genes responsible for the synthesis and assembly of MHC I molecules, as well as components involved in the processing and presentation of antigens. In addition, TNF-α pathway is upregulated in response to injury (Fig. 4 a, S4), enhancing the expression of MHC molecules and facilitating antigen presentation during inflammation.

**Fig. 4:**
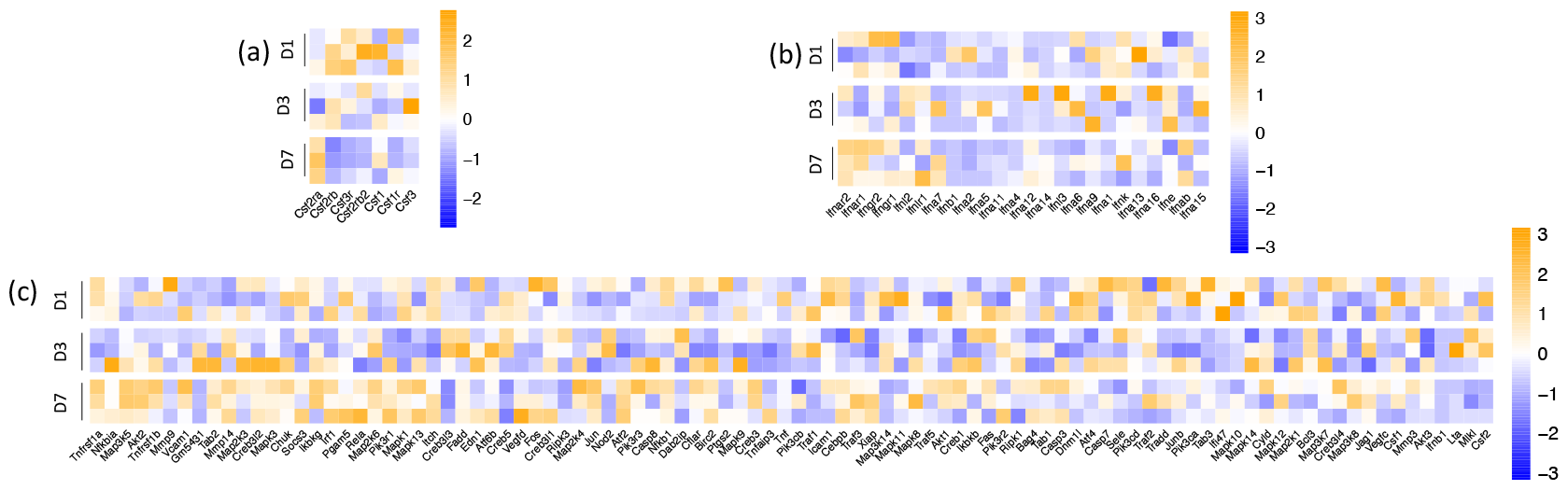
Cytokines release by Müller glia cells in response to laser injury. (a) Heatmaps of differentially expressed genes that encode receptors for CSFs in Rlbp1^GFP^ cells (Müller glia). Data are expressed as fold-changes between different time points (days 1, 3, and 7) compared to negative controls (Rlbp1^GFP^ cells from uninjured retinas). (b) Heatmaps of differentially interferon receptor-ligand genes in Rlbp1^GFP^ cells (Müller glia). Data are expressed as fold-changes between different time points (days 1, 3, and 7) compared to negative controls (Rlbp1^GFP^ cells from uninjured retinas). (c) Heatmaps of differentially expressed genes of the TNF signaling pathway in Rlbp1^GFP^ cells (Müller glia). Genes were selected from KEGG pathways (mmu04668). Data are expressed as fold-changes between different time points (days 1, 3, and 7) compared to negative controls (Rlbp1^GFP^ cells from uninjured retinas).

Our findings suggest Müller cells actively participate in retinal adaptive immunity by potentially presenting antigens to CD8^+^ T-cells. Müller glia may act as atypical APCs through interferons and TNF-α pathway.

### Glial cell-mediated antigen presentation in human retinas during degeneration

Other studies have also shown that glial cells are capable of acting APCs in preclinical models of retinal diseases; however, little is known about their role as APCs in humans.

Therefore, we first analyzed histopathological features of ocular tissue from human donor eyes by investigating the presence of drusen and structural changes in the retinal tissue (Fig. 5). Using H&E staining, we identified 15 specimens exhibiting healthy cuboidal RPE. These cells displayed a uniform shape, with centrally located nuclei and abundant melanin pigment granules in the cytoplasm, giving them a brownish appearance. The borders between adjacent RPE cells were well-defined, and the basal surface of RPE cells lay on Bruch’s membrane (Fig. 5 a). These retinal samples exhibited a normal multilayered structure with distinct cellular arrangements and were categorized as the CTRL group (Fig. 5 a, c-d). Conversely, 15 specimens showed drusen, which typically appeared as discrete, round, or oval-shaped deposits located between the RPE and Bruch’s membrane. Drusen often presented as eosinophilic (pink-staining) structures, with sizes ranging from 25 to 125 μm. Specimens with drusen exhibited a thinner retina, reduced cellular density, and structural disorganization compared to the CTRL group. This ongoing degeneration was evident from the loss of cellular components observed in the H&E-stained sections, leading us to classify these specimens as the RD group (Fig. 5 a, c-d). To further confirm the pathological state of the RD retinas compared to CTRL, we performed Picro Sirius red staining to visualize tissue fibrosis. As expected, the RD group displayed fibrotic retinas with prominent red collagen fibers, often arranged in irregular patterns or bundles. In contrast, CTRL specimens exhibited a pale pink coloration, indicating the absence of fibrosis or pathological changes characteristic of a healthy physiological state (Fig. 5 e).

**Fig. 5:**
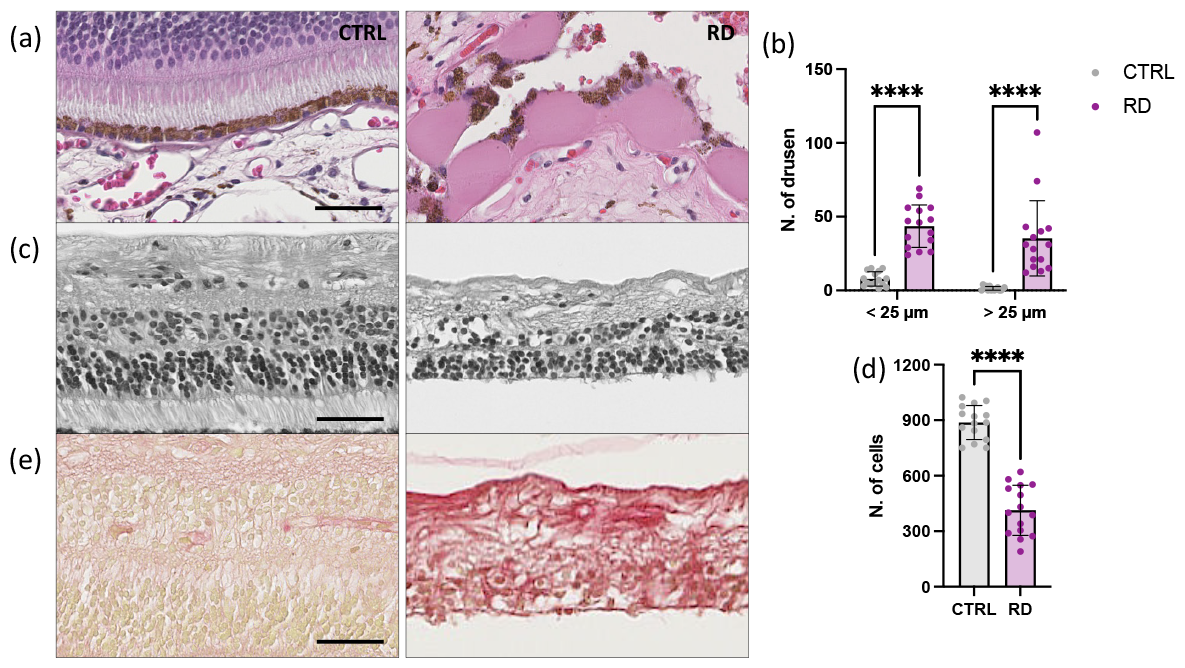
Pathological examination of ocular tissue from human donors’ eyes. (a-d) H&E staining of human retinas showing presence of drusen and its integrity. (a) Shown are representative detailed view of healthy cuboidal RPE (CTRL) and retinas presenting drusen (RD). (b) Quantification of drusen as either micro drusen (Ø < 25 μm) or hyalinized rounded deposits (Ø > 25 μm) between the RPE and the Bruch’s membrane. Significant differences (****p < 0.0001) between CTRL and RD groups were determined by using two-tailed Mann–Whitney test analysis (n = 15). (c) Shown are representative detailed view of healthy retinal structure (CTRL) and atrophic retinas (RD). (d) Quantification of cell nuclei per retina. Significant differences (****p < 0.0001) between CTRL and RD groups were determined by using two-tailed Mann–Whitney test analysis (n = 15). (e) Shown are representative sections stained with Picro Sirius red staining for histological visualization of collagen fibers and retinal fibrosis. Scale bars equals 50 μm.

Furthermore, we assessed whether resident phagocytic cells (Iba1^+^) and Müller glia (GS^+^) exhibited characteristics of APCs upon activation in the diseased retina (Fig. 6 a-d). Iba1^+^ cells were exclusively iNOS^+^ in RD specimens (Fig. 6 a-b), indicating their activation during degenerative processes and their polarization into a pro-inflammatory state. Additionally, we observed reactive Müller glia in RD samples. During retinal degeneration, over 60 % of GS^+^ cells co-expressed GFAP, compared to only 4% of GFAP^+^GS^+^ cells found in the CTRL group. The presence of GFAP^+^ Müller glia during retinal degeneration indicates their reactive and gliotic state compared to the quiescent state in healthy conditions.

**Fig. 6:**
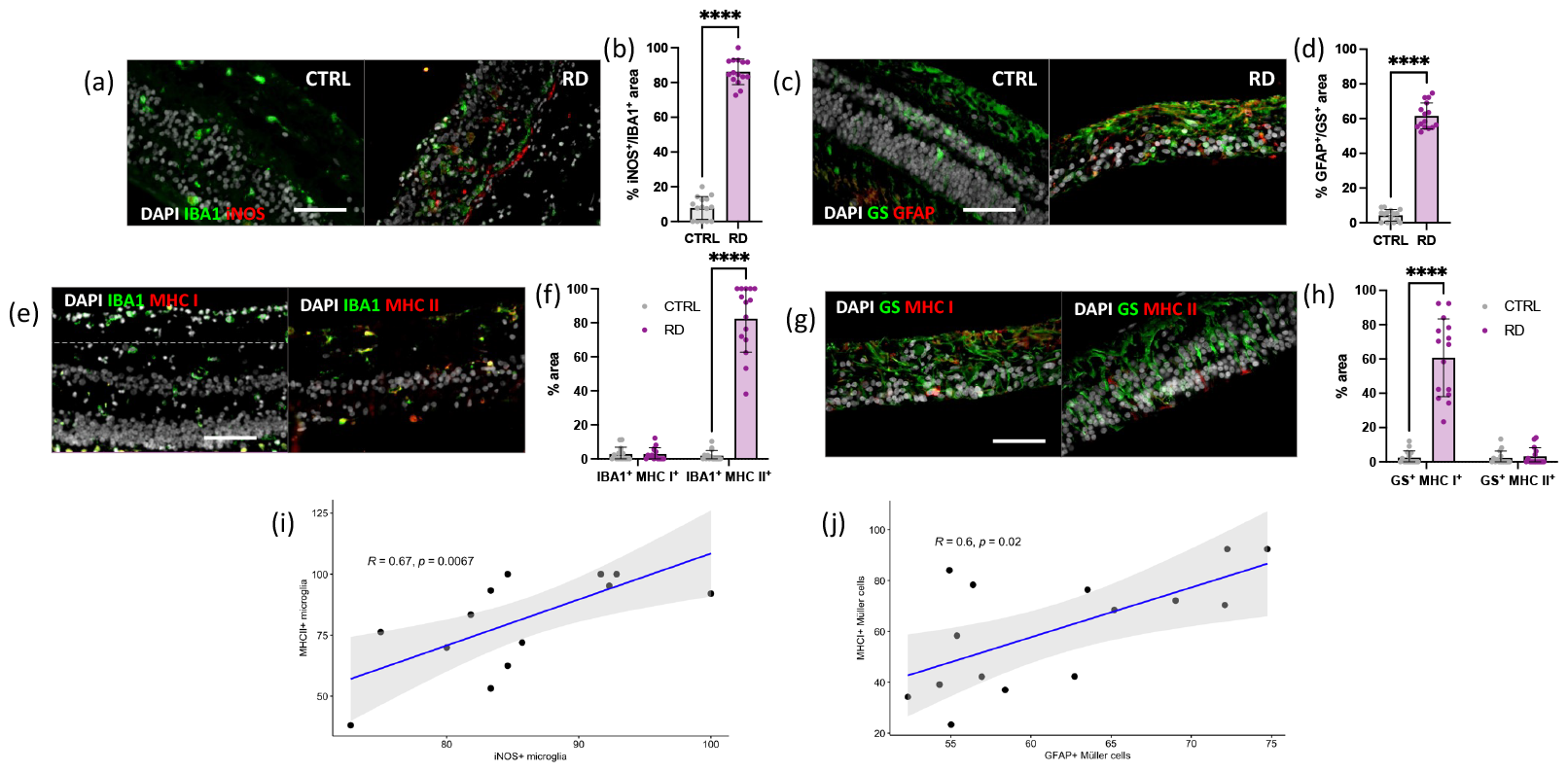
Glial responses correlate MHCs in human retina during degeneration. (a-b) Analysis of pro-inflammatory phenotype in microglia in healthy (CTRL) and degenerated retinas (RD) using a microglia marker (IBA1) and a pro-inflammatory marker (iNOS). (a) Shown are representative sections stained for Iba1 (green) and iNOS (red). (b) Quantification of the percentages of iNOS^+^IBA1^+^ area on total IBA1^+^ area in CTRL and RD retinas. Significant differences (****p < 0.0001) between CTRL and RD were determined by using two-tailed Mann–Whitney test analysis (n = 15). (c-d) Analysis of Müller cell reactivity in healthy (CTRL) and degenerated retinas (RD) using a Müller cell marker (GS) and a reactivity marker (GFAP). (c) Shown are representative sections stained for GS (green) and GFAP (red). (d) Quantification of the percentages of the GFAP^+^GS^+^ area on total GS^+^ area in CTRL and RD retinas. Significant differences (****p < 0.0001) between CTRL and RD were determined by using two-tailed Mann– Whitney test analysis (n = 15). (e-h) Analysis of MHC proteins by glia in degenerated retinas. (e) Shown are representative sections stained for Iba1 (green) and MHC I or II (red). (f) Quantification of the percentages of the MHC I^+^ IBA1^+^ and MHC II^+^ IBA1^+^ area on total IBA1^+^ area in CTRL and RD retinas. Significant differences (****p < 0.0001) between CTRL and RD were determined by using two-tailed Mann–Whitney test analysis (n = 15). (g) Shown are representative sections stained for GS (green) and MHC I or II (red). (h) Quantification of the percentages of the MHC I^+^GS^+^ and MHC II^+^GS^+^ area on total GS^+^ area in degenerated retinas. Significant differences (****p < 0.0001) between CTRL and RD were determined by using two-tailed Mann–Whitney test analysis (n = 15). Scale bars equals 100 μm. (i) Spearman correlation between MHC II^+^ IBA1^+^ cells with iNOS^+^ IBA1^+^ cells in the degenerated retinas. (j) Spearman correlation between MHC I^+^GS^+^ cells with GFAP^+^GS^+^ cells in the degenerated retinas.

In order to define the impact of retinal degeneration on glia APCs, we assessed the expression of MHC molecules in phagocytic cells and Müller glia in the retinal parenchyma comparing CTRL and RD group. Concomitant with the upregulation of iNOS in microglia and GFAP in Müller glia, we found the expression of MHC proteins only in degenerated retinas. Interestingly, different glial type expressed different classes of MHC. MHC I expression was detected in less than 3% of Iba1^+^ cells in both CTRL and RD groups. While only 1.8% of cells in CTRL specimen expressed MHC II, we observed a significant increase to 82% of phagocytic cells positive for MHC II in RD retinas. In Müller glia, an opposite trend was observed, as GS^+^ cells were negative for MHC II in CTRL and RD retinas. However, 60% of Müller glia were positive for MHC I exclusively in diseased retinas.

This *ex vivo* analysis revealed that phagocytic cells expressed iNOS and MHC I in the degenerated retina, while Müller cells showed positivity for GFAP and MHC II. Consequently, we investigated the relationship between glial reactivity and their expression of MHC molecules through correlation analysis. Statistical analysis of our data confirmed a significant positive correlation between iNOS and MHC I in Iba1^+^ cells, as well as between GFAP and MHC II in GS^+^ cells, highlighting the distinct involvement of microglia and Müller cells in the immune response within the retinal microenvironment under pathological conditions. Furthermore, our results suggest that the observed upregulation of MHC I and MHC II on microglia and Müller cells may facilitate antigen presentation to CD8^+^ and CD4^+^ T-cells, respectively, thereby modulating the adaptive immune response differently.

*In vitro* experiments were conducted to corroborate reciprocal impacts between glia and T-cells. Migration assays using transwells revealed that the subpopulation of transmigrating T-cells varied depending on the glial type. Double the number of CD4^+^ T-cells compared to CD8^+^ T-cells migrated toward microglia, while three times as many CD8^+^ T-cells compared to CD4^+^ T-cells migrated toward Müller glia (Fig 7 a-b).

**Fig. 7:**
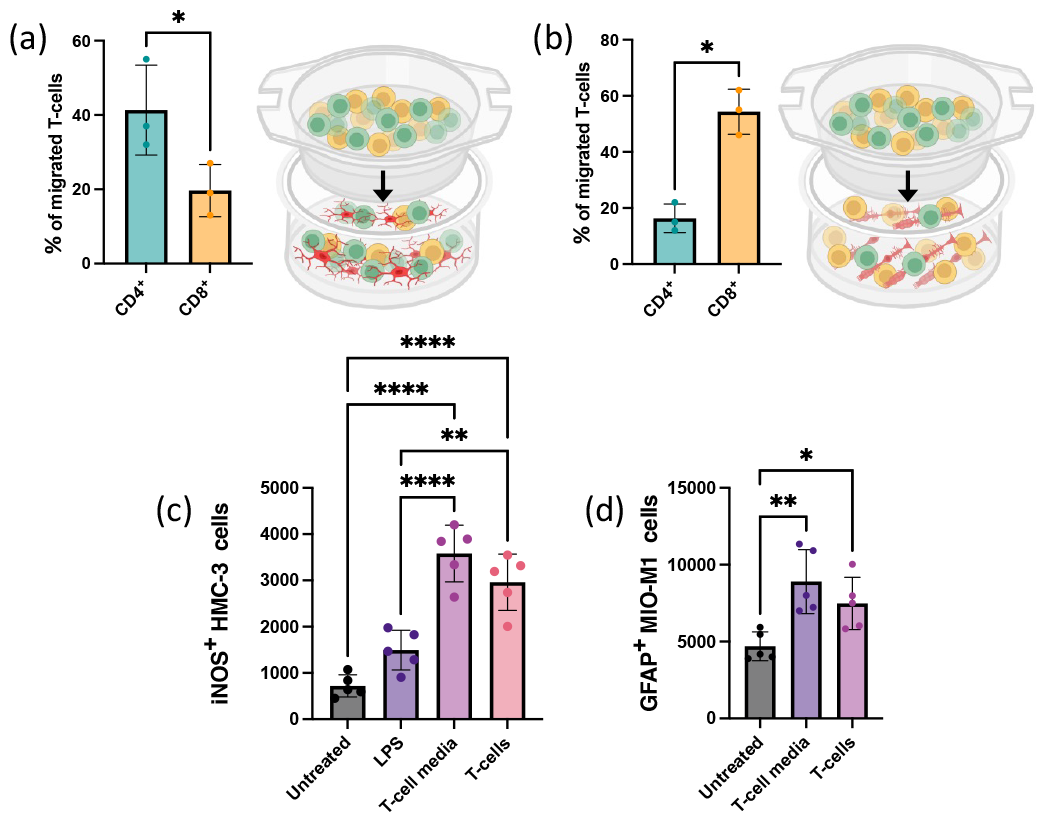
Reciprocal interplays between glia and T-cells *in vitro*. (a) Mean ± SD of the percentages of CD4^+^CD3^+^ and CD8^+^CD3^+^ cells migrated toward microglia. Significant differences (*p < 0.1) between CD4^+^CD3^+^ and CD8^+^CD3^+^ cells migrated were determined using two-tailed Mann–Whitney test analysis (n = 3). (b) Mean ± SD of the percentages of CD4^+^CD3^+^ and CD8^+^CD3^+^ cells migrated toward Müller cells. Significant differences (*p < 0.1) between CD4^+^CD3^+^ and CD8^+^CD3^+^ cells migrated were determined using two-tailed Mann–Whitney test analysis (n = 3). (c) Quantification of the iNOS^+^ HMC-3 cells untreated, treated with LPS, T-cell media or co-cultured with T-cells. Significant differences (**p < 0.01 and ****p < 0.0001) between conditions were determined by using a post hoc Bonferroni one-way ANOVA test (n = 5). (d) Quantification of the GFAP^+^ MIO-M1 cells untreated, treated with T-cell media or co-cultured with T-cells. Significant differences (*p < 0.1 and **p < 0.001) between conditions were determined by using a post hoc Bonferroni one-way ANOVA test (n = 5).

Finally, we incubated microglia and Müller cells with activated T-cells or their media to analyze their effects on glial cells. As expected, a higher number of LPS-treated microglia expressed iNOS compared to untreated cells. Interestingly, this expression increased further when incubated with activated T-cells or their media, suggesting that activated T-cells induced a pro-inflammatory phenotype in microglia (Fig. 7 c). Additionally, cultured Müller cells expressed GFAP exhibiting reactivity, but when exposed to activated T-cells or their media, this expression increased even more, confirming the impact of activated T-cells on Müller glia reactivity (Fig. 7 d).

Overall, our *ex vivo* and *in vitro* data demonstrate strong and specific T-cell-glial cell interactions. Different T-cell subpopulations are recruited by various types of glia, which in turn become more active upon interaction with T-cells.

## DISCUSSION

Since the 1980s, retinal diseases have been intricately linked to immune system involvement (Dumonde et al., 1985; Keltner et al., 1983). Numerous clinical trials have focused on modulating retinal inflammation to promote tissue regeneration, but achieving consistent treatment success has proven challenging. The underlying causes of these failures are unclear (Kim et al., 2021).

In recent years, diverse paradigms concerning retinal immunology have been challenged, including the discovery of glial cells acting as antigen-presenting cells and the role of T-cells in driving neurodegeneration (Grüntzig & Hollmann, 2019; Sutter & Crocker, 2022; Zhang & Jiang, 2023). Once retinal tissue is damaged, eye-derived antigens drain into the lymph nodes to activate T-cells, which are recruited in the retinal parenchyma. However, we still have limited knowledge about the functional role of glial cells as APCs within the damaged retinal tissue (Schetters et al., 2017). We used a laser coagulator to induce photoreceptor disorganization, glial proliferation and scarring common macroscopic features of human retinal degenerative diseases [e.g., AMD; (Conedera et al., 2021)]. Previous studies have shown that laser-induced injury triggers the migration of resident immune cells and macroglial reactivity specifically within the site of injury (Chidlow et al., 2013; Miller et al., 2019). Therefore, the laser-induced injury model proves invaluable for investigating glial biological processes directly within the damaged area, while using adjacent tissue as a control.

Activated microglia and resident macrophages are frequently observed in various neuropathologies and implicated in their progression (Goddery et al., 2021). Intriguingly, many neurological conditions characterized by their activation also exhibit evidence of infiltrating T-cell populations (Behnke et al., 2020). As microglia are known to have the capability to present antigens to T-cells, these observations have spurred further investigation into the potential role of microglia as antigen-presenting cells in several retinal pathologies (Biswas, 2023). In line with the literature, our data from immunofluorescence and RNAseq confirmed the ability of resident immune cells to act as APCs upon laser-induced injury. We observed an upregulation of genes and proteins related to the MHC II pathway, unlike MHC I expression, which was not significantly different from the baseline. Among the upregulated considerably molecules, we found H2-Eb2, Cd74 and Ciita, which are important in aiding in peptide presentation to CD4^+^ T-cells, stabilizing MHC II molecules, and regulating MHC II gene transcription, respectively. These data showed that resident immune cells upregulated MHC II and its pathway, suggesting their ability to present antigens to CD4^+^ helper T-cells. Supporting our results, similar interactions between MHC II^+^ resident immune cells and CD4^+^ T-cells have been documented in other contexts, such as Tau-mediated neurodegeneration and aging in the murine brain (Askin & Wegmann, 2023; Kellogg et al., 2023; Zhang et al., 2022). In the retina, their interplay is well-characterized only in animal models of autoimmune uveitis (Lipski et al., 2017; Okunuki et al., 2019) and little is known in retinal degenerative diseases. However, we know that CD4^+^ T-cells are recruited to the parenchyma in mice with experimental ischemic retinopathy and laser-injuries and both models also involved glial activation (Conedera et al., 2023; Khanh Vu et al., 2020; Wagner et al., 2021). Interestingly, other recent publications implied the connection between resident immune cells and CD4^+^ T-cells showing that the depletion of CD4^+^ T-cells leads to the accumulation of pro-inflammatory microglia (Llorián-Salvador et al., 2024). Thus, resident immune cells can stimulate CD4^+^ T-cell migration to the damaged retina as well as T-cells can affect immune cell activation once they interplay with T-cells. We indeed detected a significant upregulation of Csf1r compared to baseline, which suggests an increased demand for CSF signaling. This data implies heightened responsiveness of immune cells to CSF1, the ligand for CSF1R, leading to their activation and proliferation. This activation enhances immune cells’ ability to respond to injury signals in the retina, potentially prolonging their reactive state.

While microglia are recognized as primary antigen-presenting cells in the retina, it’s noteworthy that Müller cells, the principal glial cells of the retina, can also exhibit antigen-presenting capabilities. This occurs in response to inflammatory conditions triggered by various pathological stimuli such as infection, trauma, or autoimmune reactions. Under such circumstances, Müller cells upregulate antigen-presenting molecules and actively participate in presenting antigens to T-cells. This capability allows Müller cells to interact with the immune system and modulate immune responses within the retina. Our results confirm that Müller cells possess the machinery required for antigen presentation, as evidenced by the expression of MHC I protein. Contrarily, MHC II protein level in Müller cells remained comparable with basal levels throughout the injury response. Müller cells contribute to the injury response by acting as atypical APCs 7 days post injury, unlike resident immune cells that acquired the capacity to present antigens to T-cells already on day 3. The timing of microglial and macroglial responses to injury varies due to differences in their cellular properties, functions, and activation mechanisms. As sentinel cells, microglia are poised to respond rapidly to injury signals due to their constant surveillance and activation state. Microglia undergo rapid activation and migration to the injury site upon detecting injury-associated signals, such as damage-associated molecular patterns (DAMPs) or cytokines released by damaged cells. While Müller cells also respond to injury, their primary function is maintaining retinal homeostasis, providing metabolic support to neurons, and regulating extracellular ion and neurotransmitter levels. Furthermore, their delayed response may be related to the need for sustained changes in the retinal microenvironment, such as prolonged inflammation or cellular damage, which can activate Müller cells and induce their participation in forming a gliotic scar. Transcriptome analysis confirmed that Müller cells predominantly upregulated genes related to antigen processing 7 days after injury, coinciding with the expression of MHC I protein (Kelly & Trowsdale, 2019). Psme1, various Tap genes, and Canx were significantly upregulated in Müller cells, demonstrating their ability to generate peptides for MHC class I presentation, facilitate peptide transport for MHC class I loading, and aid in the folding and assembly of MHC class I molecules. Thus, these data demonstrate that Müller cells upregulate MHC class I and its pathway, suggesting their capacity to present antigens to CD8^+^ cytotoxic T-cells. This differs from resident immune cells, which typically present antigens to CD4^+^ T-cells during injury response in the retina. Although Müller cells are involved in virtually every retinal disease, their role in neuroinflammation is still poorly understood. Our results obtained from a preclinical model of retinal degeneration are supported by previous studies *in vitro* (Lorenz et al., 2021; Schmalen et al., 2021). They used proteomic analysis to demonstrate that Müller cells express both MHC molecules under different pro-inflammatory stimuli. Similar to Müller cells, astrocytes can also express both MHC I and II during neurodegeneration. Infiltrated CD4^+^ T-cells were observed interacting with MHC II-expressing astrocytes in pathological conditions such as Parkinson’s disease and multiple sclerosis (Cornet et al., 2000; Rostami et al., 2020). *In vitro* studies further revealed that astrocytes sustained antigen-specific CD4^+^ T-cell proliferation only in the presence of IFNγ, facilitated by the astrocytic expression of MHC II (Cornet et al., 2000). However, examination of the distribution of astrocytic MHC-positive cells in relation to lesion architecture revealed distinct patterns: MHC II^+^ cells were confined exclusively to the lesion edge, while MHC I^+^ astrocytes were widely present at both the lesion edge and in the gliotic lesion center (Ransohoff & Estes, 1991). These findings suggest an association between gliosis and MHC I expression, a relationship further supported by *in vitro* analysis (Bombeiro et al., 2017). Indeed, previous publications using the same injury model and laser setup indeed demonstrated reactive gliosis on day 7, coinciding with our observation of MHC I pathway activation. This supports the notion that the expression of MHC molecules varies depending on the reactive status of Müller cells. However, further analyses are required to establish the direct association between gliosis and the capacity of Müller cells to function as APCs.

Glia and T-cells interact with each other through the secretion of certain cytokines and pro-inflammatory molecules, which have significant effects on the expression of MHC molecules (Johansson et al., 2008; Raval et al., 1998). We indeed found an upregulation of Csf2ra in reactive Müller cells providing additional evidence of their potential role in antigen presentation via MHC I molecules, contributing to immune surveillance and responses within the retina (Saita et al., 2022). Additionally, we found significant differences in IFN expressions and the TNF-α pathway, which are closely linked to the regulation of MHC molecule expression and antigen presentation (Cantaert et al., 2010; Silginer et al., 2017). Previous publications showed that IFN and TNF-α induces the expression of major histocompatibility antigens by human retinal glial cells (Chen et al., 2022; Heng et al., 2019; Mano et al., 1991). Additionally, both cytokines TNF-aggravated gliosis in the retina and brain (Hu et al., 2020; Wangler et al., 2022; Yong et al., 1991).

Inflammation and gliosis play harmful roles in retinal degeneration in mice, as observed in humans (Bora et al., 2003); however, immune and glial responses can differ between the two species (Bjornson-Hooper et al., 2022). These differences rely on variations in the extent, timing, and specific cellular mechanisms. Hence, a thorough understanding of the involvement of glia in the immune response during the progression of retinal degeneration in humans is crucial for developing targeted therapies. To achieve this, we conducted retrospective studies using formalin-fixed and paraffin-embedded specimens from donor eyes post-cornea removal.

Differences in antigen presentation by microglia and macroglia following laser-induced injury in the mouse retina were confirmed in human retinas. Specimens with drusen were characterized as degenerative (RD). They exhibited features such as a thinner retina, reduced cellular density, structural disorganization, and prominent fibrotic tissue with collagen fibers. Interestingly, we found a correlation between MHC expression and glial response. Most resident immune cells (Iba1^+^) appeared activated (iNOS1^+^), expressing MHC II but not MHC class I. In contrast, Müller cells were also reactive (GFAP+) and expressed only MHC class I but not MHC class II. These results suggest a role in regulating the specific response of CD8^+^ and CD4^+^ T-cells by the expression of MHC class I and class II molecules, respectively (Sprent et al., 1990). Thus, activated glial cells can influence the nature and extent of the immune response within the retina, potentially promoting either a pro-inflammatory or anti-inflammatory environment and thus affecting the progression of degeneration (Terrabuio et al., 2023). To further confirm this, we conducted in vitro experiments assessing the interplay between different glial and T-cell types. The subpopulation of T-cells that migrated towards the glial population depended on their cell type: CD4^+^ T-cells responded significantly stronger to LPS-activated microglia than CD8^+^ T-cells. Conversely, the latter migrated preferentially towards cultured Müller cells rather than microglia. Furthermore, directed migration depended on soluble factors, and similar results were obtained using supernatants collected from the respective glial population. These findings suggest optimized mechanisms for both activation-stage and environment-specific attributes (Krummel et al., 2016).

In summary, retinal glial cells play an active role in modulating the immune response following insults to the retinal parenchyma. Therefore, understanding the dynamics of glia-mediated antigen presentation in the retina during degeneration is crucial for developing targeted therapeutic strategies for retinal diseases. By modulating the immune response orchestrated by glial cells towards a more beneficial one, it may be possible to mitigate retinal degeneration and even potentially restore vision in affected individuals.

## SUPPLEMENTARY FIGURES

**Fig. S1:**
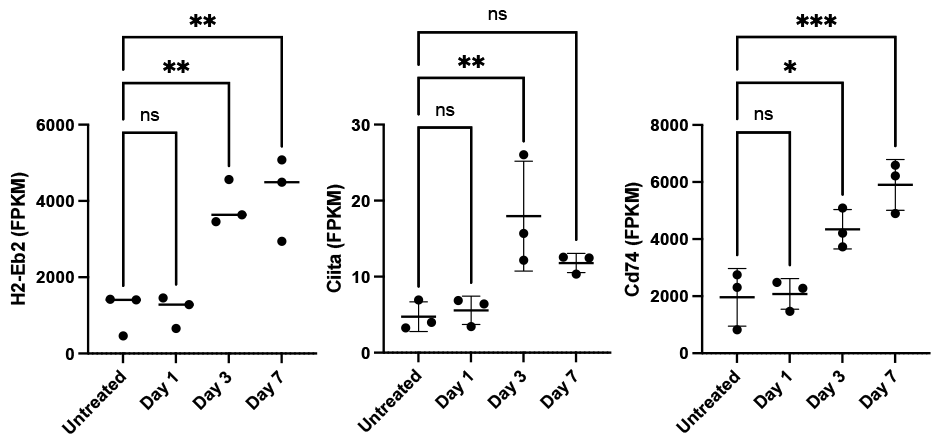
Gene expression level of molecules playing critical roles in the MHC class II pathway. RNAseq data quantifying expression of RNAs encoding H2-Eb2, Ciita and Cd74 as fragments per kilobase of transcript per million mapped reads (FPKM).

**Fig. S2:**
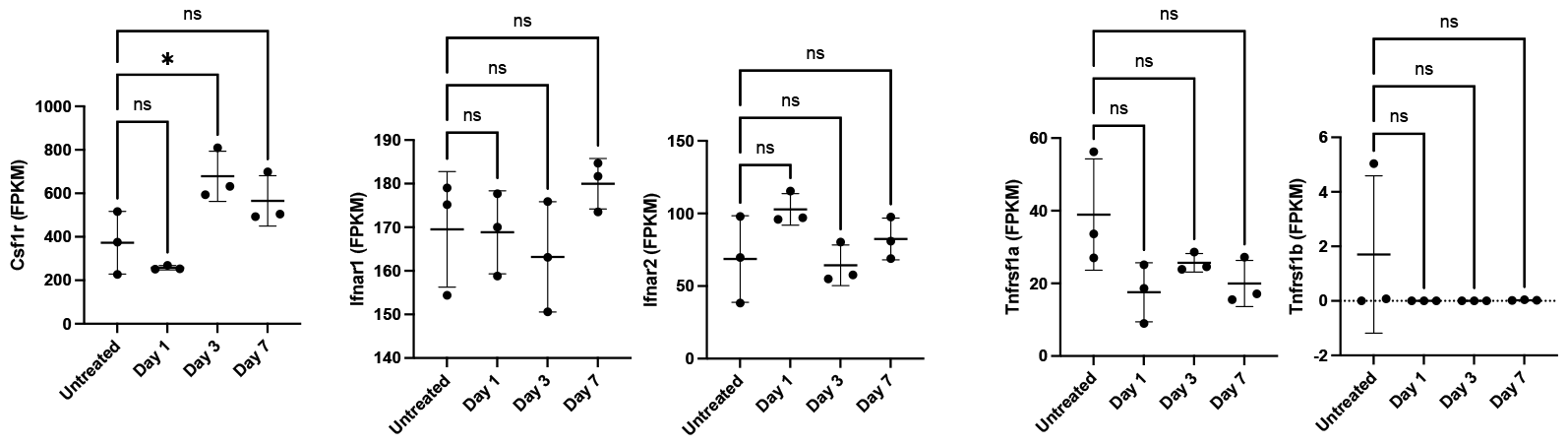
Expression level of Csf genes, interferons and molecules playing critical roles in the TNF signaling. RNAseq data quantifying expression of RNAs encoding Csf1r, Ifnar1, Ifnar2, Tnfrsf1a and Tnfrsf1b as fragments per kilobase of transcript per million mapped reads (FPKM).

**Fig. S3:**
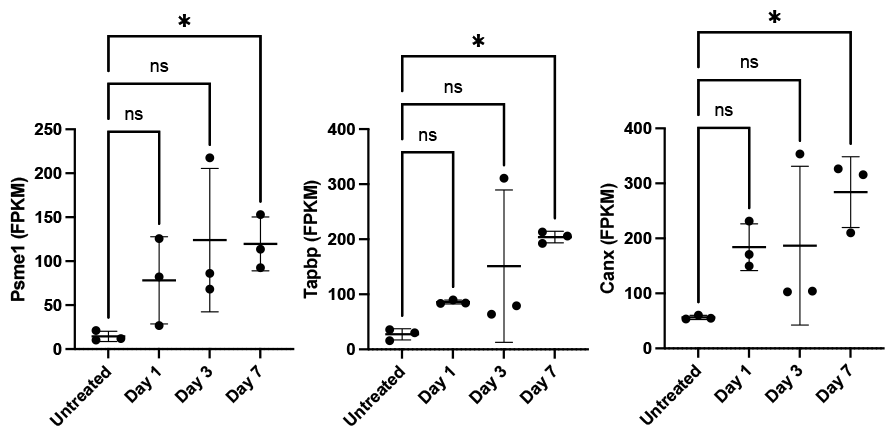
Gene expression level of molecules playing critical roles in the MHC class I pathway. RNAseq data quantifying expression of RNAs encoding Psme1, Tapbp and Canx as fragments per kilobase of transcript per million mapped reads (FPKM).

**Fig. S4:**
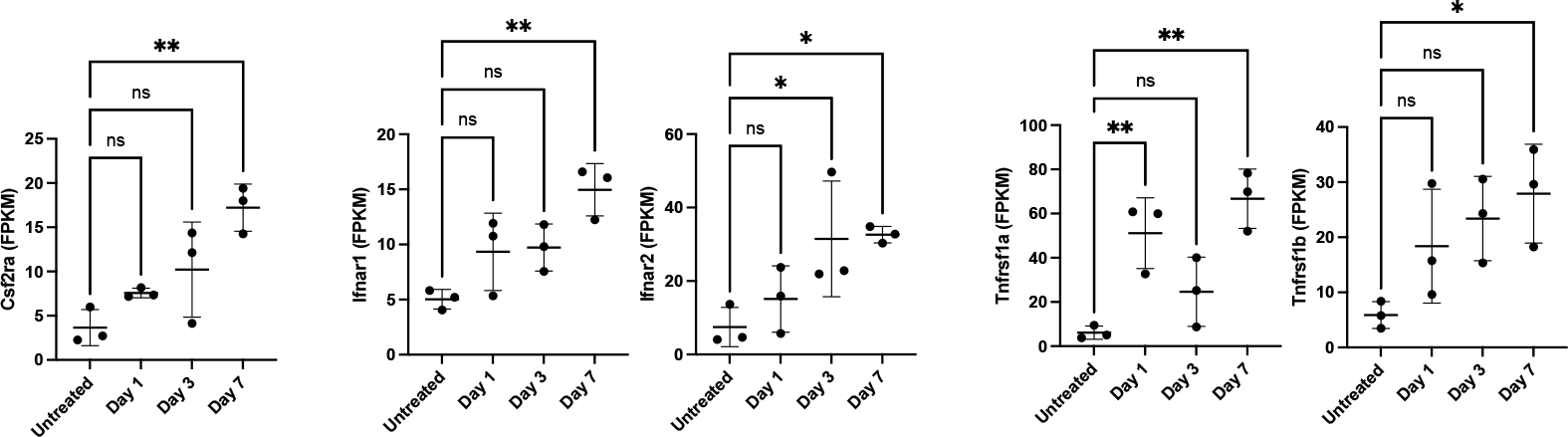
Expression level of Csf genes, interferons and molecules playing critical roles in the TNF signaling. RNAseq data quantifying expression of RNAs encoding Csf2ra, Ifnar1, Ifnar2, Tnfrsf1a and Tnfrsf1b as fragments per kilobase of transcript per million mapped reads (FPKM).

**Fig. S5:**
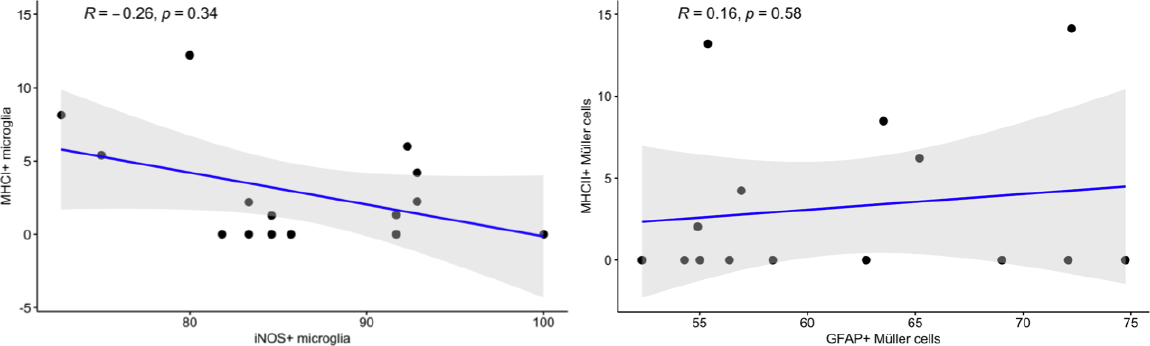
Relationship between glial reactivity and their expression of MHC molecules. Spearman correlation between MHC I and iNOS^+^Iba1^+^ cells and MHC II with GFAP^+^GS^+^ cells.

